## Supplemental Material for "Functional and structural characterization of interactions between opposite subunits in HCN pacemaker channels"

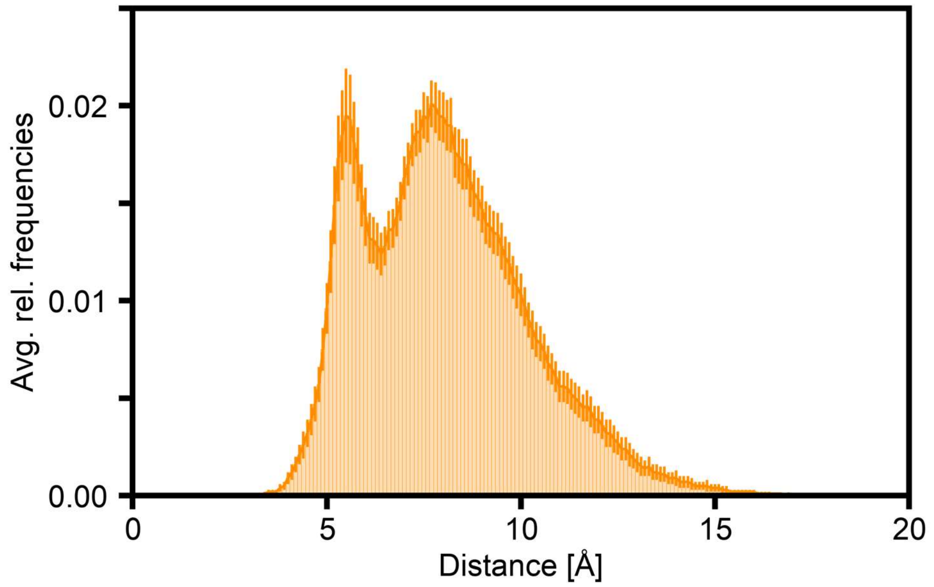


**Figure S1: Distance measurements between K464E and M155.**
Histogram (bin size 0.1 Å) of the minimal distance between the side-chain oxygen atom in residue K464E and any backbone atom in residue M155. The histogram is normalized to the sum of all bins. The average values were calculated over all four subunits of the mHCN2 channel and over 20 independent replicas (*n* = 80). The error bars denote the standard error of the mean.

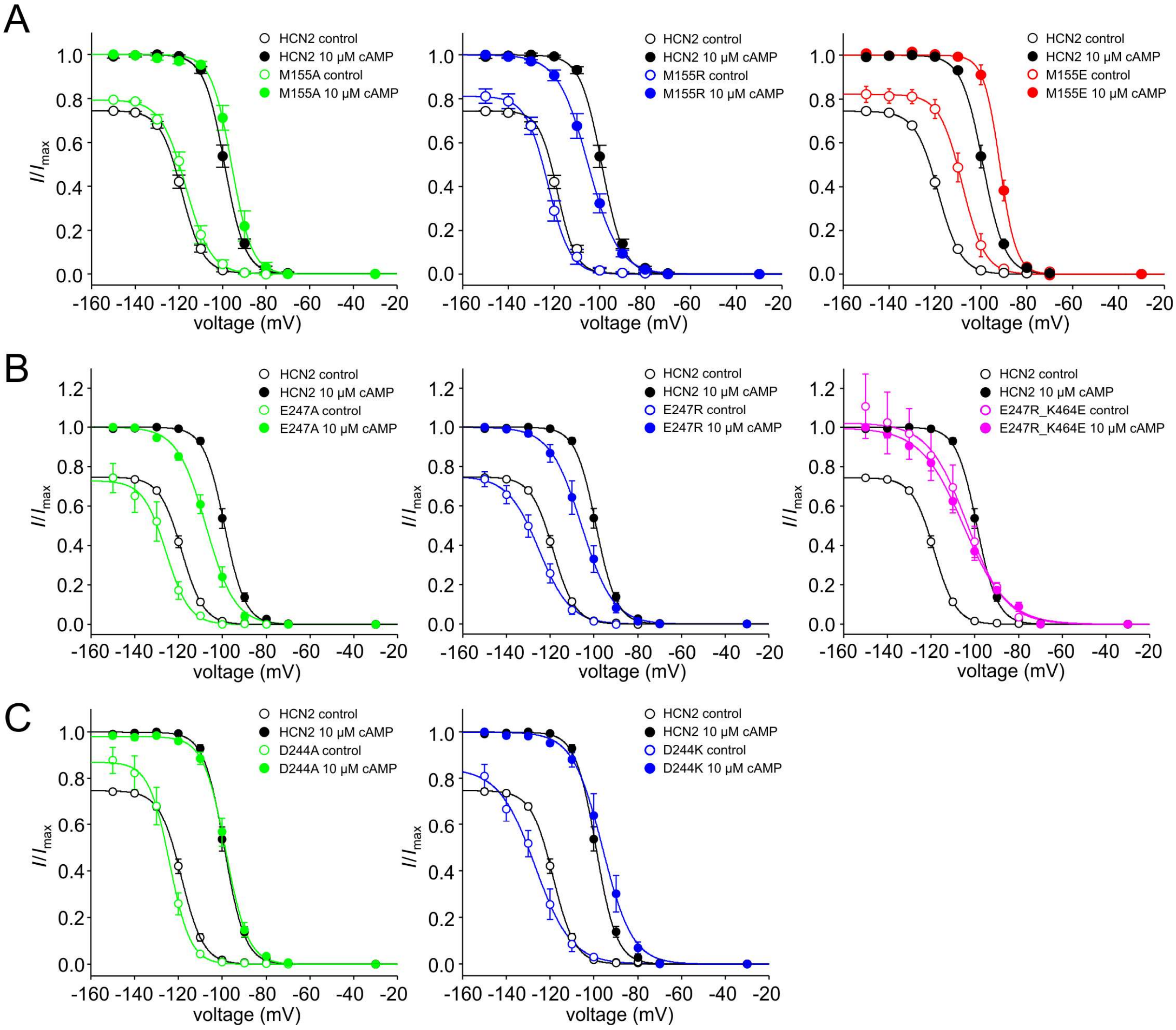


**Figure S2: Steady-state activation relationships for mutated constructs in comparison with mHCN2 wildtype.**

Shown are steady-state activation relationships for A) M155A, M155R, M155E, B) E247A, E237R, E247R_K464E, C) D244A, D244K in the absence and presence of saturating [cAMP] (colored symbols as incidicated). In each case, open and filled symbols represent mHCN2 in the absence and presence of saturating [cAMP], respectively. The Boltzman equation was fitted to the data yielding $V_{1/2}$ and $z\delta$.

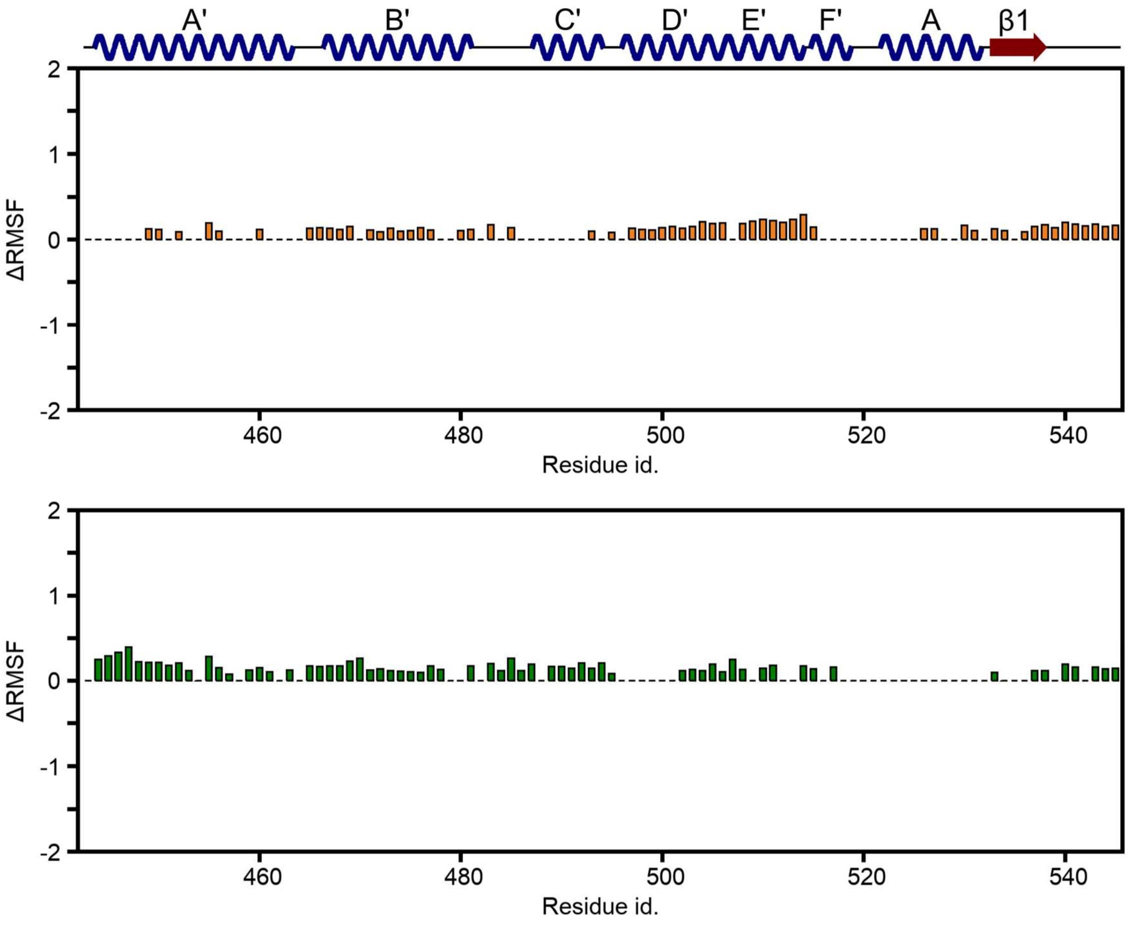
A'
B'
C'
D'
E'
F'
A
β1
ΔRMSF
2
1
0
-1
-2
460
480
500
520
540
Residue id.
ΔRMSF
2
1
0
-1
-2
460
480
500
520
540
Residue id.

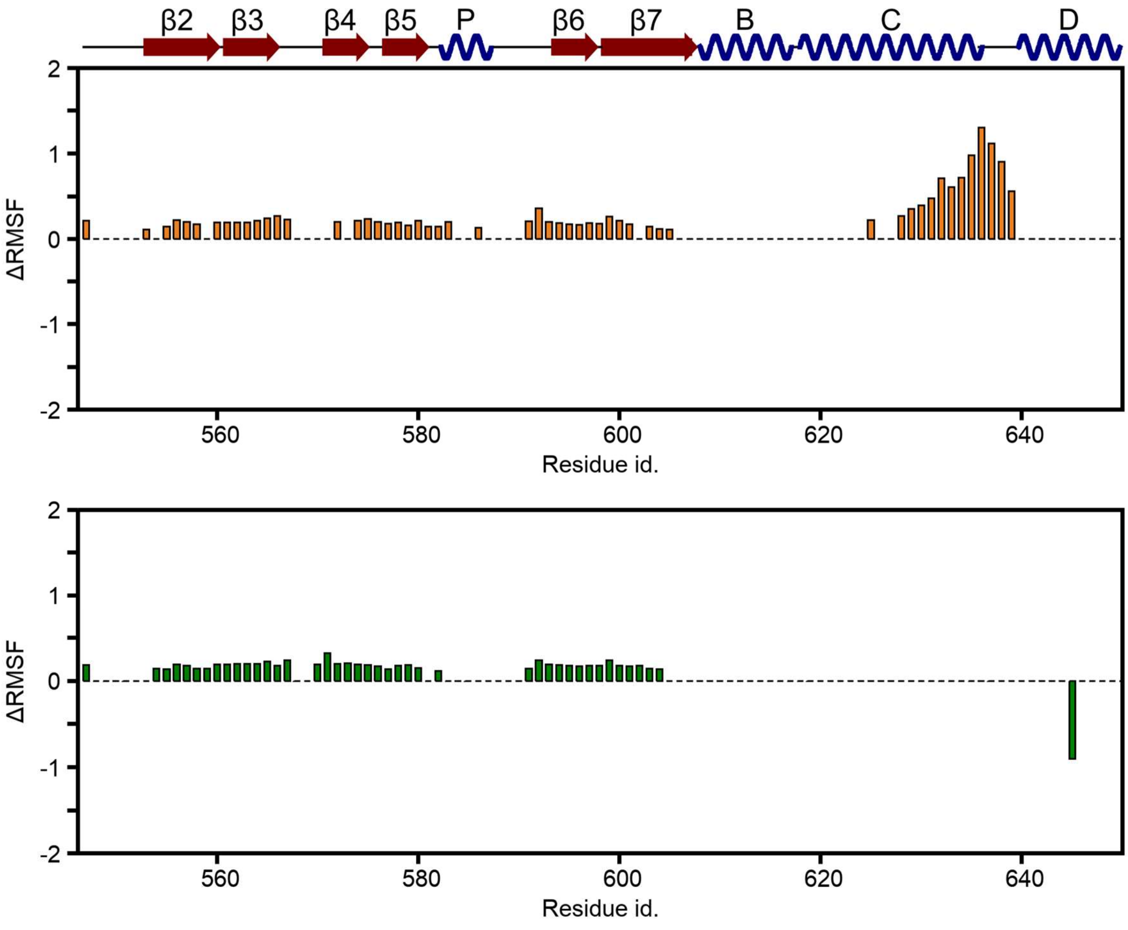
β2
β3
β4
β5
P
β6
β7
B
C
D
ΔRMSF
2
1
0
-1
-2
560
580
600
620
640
Residue id.
ΔRMSF
2
1
0
-1
-2
560
580
600
620
640
Residue id.

**Figure S3: Changes in side-chain mobility within the CL-CNBD induced by cAMP binding or K464E substitution.**

The bar plots show the residue-wise average ΔRMSF (root mean square fluctuations; see also eq. (4); $n$ = 80 independent replicas)). The orange bars show the ΔRMSF for the cAMP-bound wild type channel, and the green bars the ΔRMSF for the *apo* K464E channel, with respect to the *apo* wild type channel. If the residue-wise RMSF is not significantly different to the *apo* wild type channel (in the case of $p > 0.05$; $p$ value by $t$ test), ΔRMSF was set to zero. The top two panels show the ΔRMSF for residues 443 – 546 and the lower two panels for residues 547 – 650. The secondary structure of the CL-CNBD is schematically visualized, with helices shown as blue springs, β-strands as red arrows, and loops as black lines. The nomenclature was adapted from ref. Lee and MacKinnon (2017) (1).

A

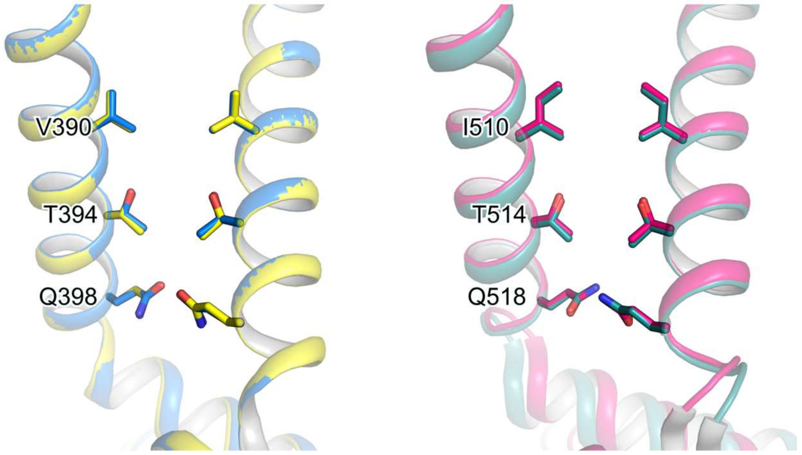


B

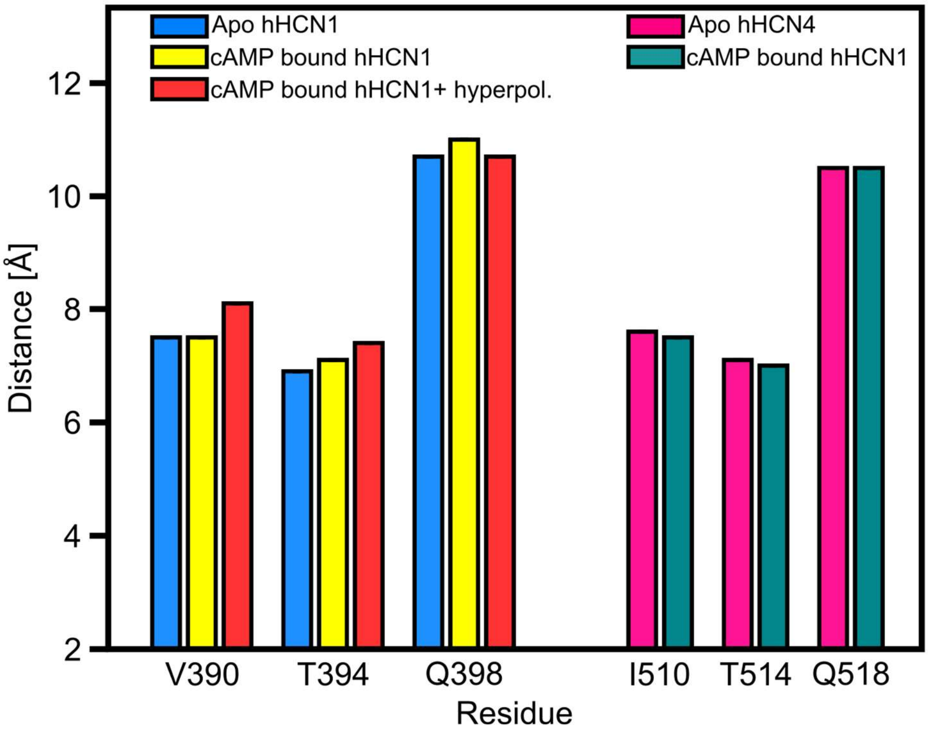


**Figure S4: Conformational analyses of the channel gate of experimental structures.**
**A:** The left structure shows the superposition of the *apo* (light blue, PDB 5U6O (Lee and MacKinnon, 2017)) and cAMP-bound (yellow, PDB 5U6P (Lee and MacKinnon, 2017)) gate region of the hHCN1, and the right panel the *apo* (magenta, PDB 6GYN (Shintre et al., to be published)) and cAMP-bound (dark cyan, PDB 6GYO (Shintre et al., to be published) hHCN4 channel. The protein structure is always shown as cartoon representation with amino acids forming the gate shown as stick models. **B:** Distance measurements between the $C_{\beta}$-atoms of two opposite amino acids that form the gate. The bar plot depicts single measures based on the available 3D structures (1-3).

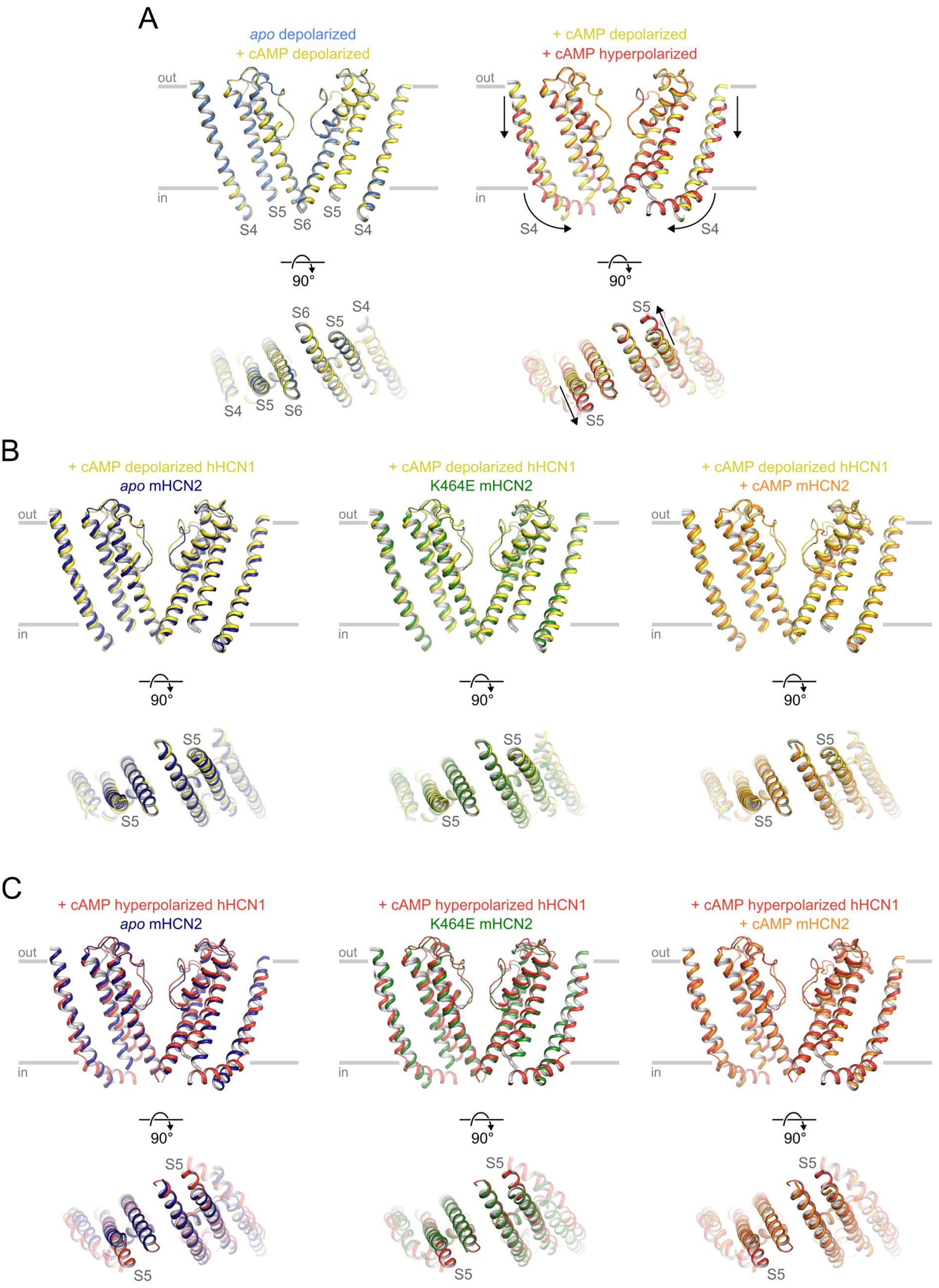


**Figure S5: Comparative analyses of the S4/S5/S6 transmembrane portion in HCN channels.**
A: Superposition of the S4/S5/S6 transmembrane portion of the hHCN1 with only two opposite subunits shown. The left panel shows the superposition of the apo (blue) and cAMP bound (yellow) structures in the depolarized conformation (1). The right panel shows the superposition of the depolarized (yellow) and hyperpolarized (red) conformations bound to cAMP (1, 4). In the right panel, structural changes induced by hyperpolarization (lowering

and bending of the S4 helix and movement of the S5 helix) are visualized by arrows. In the lower panels, the channels are rotated by 90°, such that the intracellular portion is now oriented towards the reader. The gray labels depict relevant structural domains (the S4, S5, and S6 helices). The gray bars depict the approximate location of the membrane bilayer. The layout is fully applicable to panels B and C. B, C: From left to right, the panels show the superposition of the average apo wild type mHCN2 (dark blue), the average K464E mHCN2 (green), and the average cAMP bound wild type mHCN2 (orange) channels from MD simulations onto the cAMP bound and depolarized conformation (B) or cAMP bound and hyperpolarized conformation of hHCN1 (4).

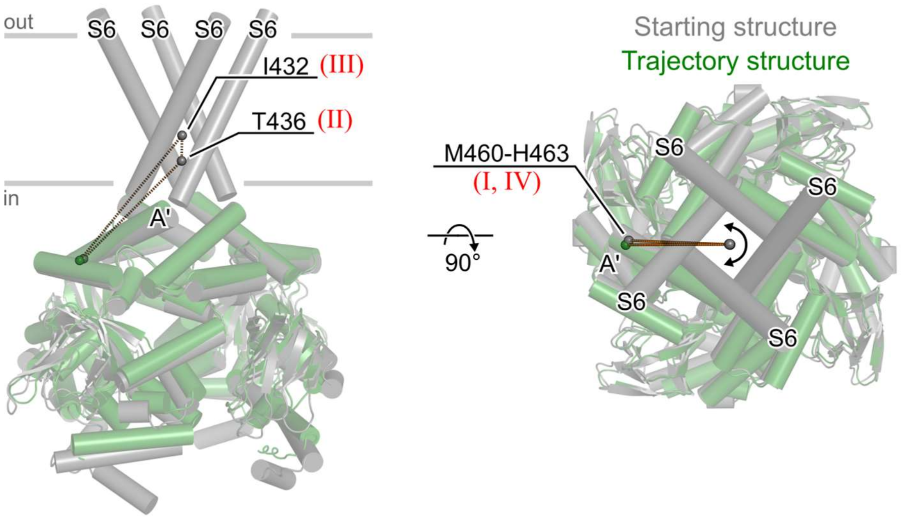


**Figure S6: Schematic definition of the rotation angle measurements.**
To investigate the rotation of the CL-CNBD relative to the channel pore of the starting structure, we measured the dihedral angle defined by the four reference points **I** – **IV** (red labels): **I**) The center of mass (COM) of $C_\alpha$-atoms of the four-terminal residues (M460-H463) of the A'-helix of the C-linker of the starting structure (gray cartoon representation), **II**) the COM of $C_\alpha$-atoms of T436 on the S6 helix of each subunit of the starting structure, **III**) the COM of $C_\alpha$-atoms of I432 on the S6 helix of each subunit of the starting structure, and **IV**) the COM of $C_\alpha$-atoms of the four-terminal residues (M460-H463) of the A'-helix of the C-linker, but now considering the atoms of structures throughout the MD trajectory (green cartoon representation). The COMs of **I** – **IV** are depicted as spheres, and the dihedral is schematically represented by dashed lines. The left structure panel shows the channel from the side, the right from the top (the extracellular domains are now oriented towards the viewer). The gray bars depict the approximate location of the membrane bilayer. Only the S6 helices of the channel pore are shown for clarity. Helical structures are shown as cylinders.
